# Spatial spread of white-nose syndrome in North America, 2006-2018

**DOI:** 10.1101/2021.01.28.428526

**Authors:** Andrew M. Kramer, Alex Mercier, Sean Maher, Yaw Kumi-Ansu, Sarah Bowden, John M. Drake

**Affiliations:** Department of Integrative Biology, University of South Florida, Tampa, FL, 33620; Department of Biology, Missouri State University, Springfield MO 65897; Odum School of Ecology, University of Georgia, Athens GA 30602; Kapili Services, LLC, Orlando FL 32826; Center for Ecology of Infectious Diseases, University of Georgia, Athens, GA 30602

**Keywords:** *Pseudogymnoascus destructans*, spatial dynamics of disease, network model

## Abstract

White-nose syndrome has caused massive mortality in multiple bat species and spread across much of North America, making it one of the most destructive wildlife diseases on record. This has also resulted in it being one of the most well-documented wildlife disease outbreaks, making it possible to look for changes in the pattern of spatial spread over time. We fit a series of spatial interaction models to the United States county-level observations of the pathogenic fungus, *Pseudogymnoascus destructans*, that causes white-nose syndrome. Models included the distance between caves, cave abundance, measures of winter length and winter onset, and species richness of all bats and hibernating bats only. We found that the best supported models included all of these factors, but that the particular structure and most informative covariates changed over the course of the outbreak, with winter length displacing winter onset as the most informative measure of winter conditions, and evidence for the effects total species richness and hibernation varying from year to year. We also found that weather had detectable effects on spread. While the effect sizes for cave abundance and species richness were relatively stable over the length of the outbreak, distance became less important as time went on. These findings indicate that although models produced early in the outbreak captured important and consistent aspects of the spatial spread of white-nose syndrome, there were also changes over time in the factors associated with spread, suggesting that forecasts may be improved by iterative model refinement.

## Introduction

The outbreak of white-nose syndrome (WNS) in North American bats is driving massive declines in multiple hibernating bat species (Frick et al., 2010, 2015; Langwig et al., 2012, Thogmartin et al., 2012) and continues to spread across the continent (Lorch et al., 2016; Hoyt et al., 2021). The etiological agent of WNS is the psychrophilic fungus *Pseudogynmoascus destructans* (Lorch et al., 2011; Verant et al., 2012; Warnecke et al., 2012), which is known to infect at least 12 species of North American cave-dwelling bats (Turner et al., 2014; Frick et al., 2015, 2016; Hoyt et al., 2021), and has caused declines of up to 99% in abundance of *Myotis septrionalis*, *M. lucifugus*, *M. sodalis* and *Perimyotis subflavus* populations (Frick et al., 2010, 2015; Langwig et al., 2012; Thogmartin et al., 2012). Since its earliest detection in Schoharie County, NY in 2006, the pathogen has been detected in hibernacula, places including caves and mines where bats hibernate, in 39 states (Hoyt et al., 2021). This combination of host mortality and spatial extent makes WNS one of the most damaging wildlife diseases known, along with chytridiomycosis in frogs (Rohr et al., 2008) and facial tumor disease in Tasmanian devils (McCallum et al., 2007).

The well-documented spread of WNS since 2006 offers a valuable opportunity to consider how our understanding of the spatial dynamics of this disease has changed over time. A comparison of multiple models fit to the pattern of spatial spread during the first 6 years of WNS expansion in the United States found that a gravity model based on distance between counties and the number of hibernacula in each county better explained the observed pattern of spatial spread than models based solely on distance (Maher et al., 2012). The best-supported model also included a covariate for the average winter length in each county whereby longer winters were associated with a greater probability of occurrence of *P. destructans* (Maher et al., 2012). In addition to determining the role of distance, hibernacula abundance, and climate in the observed spread of WNS, this model was used to create forecasts for the future spread of WNS, summarized as the median number of new counties infected each year and the distance from origin over time (Maher et al., 2012). These forecasts also captured the order in which counties recorded infections in many cases (USFWS; www.fws.gov/whitenosesyndrome/maps.html, 2019).

*Pseudogynmoascus destructans* has continued to spread steadily across North America, providing an ever-lengthening time series of newly infected counties. (It appears that once a county has been infected, it never becomes “uninfected” again.) Studies of the disease dynamics within and between hibernacula have highlighted the role of environmental conditions within hibernacula (Wilder et al., 2011; Langwig et al., 2012; Lilley et al., 2018), finding that longer time spent hibernating leads to higher fungal loads (Langwig et al., 2020) and mortality (Lorch et al., 2011, Warnecke et al., 2012). Ongoing research also aims to determine whether autumn swarming or winter movements are more important to transmission between hibernacula (Kramer et al., 2019; Langwig et al., 2015, 2020). In this context, the previous finding that winter duration increases spread rate raises the question of whether the average length of winter (as tested by Maher et al. in 2012) matters because of regional differences in bat behavior or whether the specific conditions of each year drive spread. If the latter is the case, differences in spread rate could correlate with the number of hibernacula visited during autumn swarming or outcomes of mid-winter movements. The larger number of winter observations, compared to when the first model was created, offers additional information for answering this question.

Spread since 2012 also included long-distance dispersal to Washington in the winter of 2015-2016 (Lorch et al., 2016). The Maher et al. (2012) model predicted a median year of arrival in this region of 2026 based on a probabilistic forecast with a wide range of possible arrival times. This raises the question whether the long distance spread to Washington was an event consistent with the existing understanding of spatial dynamics of spread. Or did the short period of disease progression to that point prevent effective forecasts of future dynamics? While the dispersal of *P. destructans* to Washington has been suggested to be human-mediated (Hoyt et al., 2021), human-aided dispersal may have been a factor in earlier spread as well, and therefore implicitly recognized by the model.

Here, we revisit the spatial spread of WNS to assess how well earlier models matched subsequent spatial dynamics and factors influencing disease transmission. We extend the models previously developed to focus on three specific questions: 1) Do year-to-year weather conditions have a detectable relationship to spatial spread? 2) Was bat species richness, which was not well supported in the earlier modeling approach, a more important factor in subsequent spread? and 3) Were the effects of various factors in the original model stable during subsequent spread in space and time?

## Methods

### Data

Time of infection by county was obtained from the U.S. Fish and Wildlife Service (USFWS; www.fws.gov/whitenosesyndrome/maps.html, 2019), recording the first year in which either WNS was observed, the infectious agent *P. destructans* was discovered (e.g. in a soil sample), or both. Given that fall and winter are the crucial periods for transmission (Langwig et al., 2020; Hoyt et al., 2021) and detection (Warnecke et al., 2012; Langwig et al., 2015), the epidemic year was assumed to begin June 1^st^ and end May 31^st^.

Number of caves in each county in the contiguous United States was provided by Maher et al. (2012) and the Euclidean distance between county centroids from the same reference (Maher et al., 2012). County-level bat species richness was estimated from NatureServe (www.natureserve.com) for the 46 bat species occurring in the contiguous United States at any point in the year, as provided by Maher et al. (2012). Each species was classified as hibernating or non-hibernating based on USGS information (USGS, 2020).

We quantified average and annual winter conditions using the minimum daily temperature from all operational weather stations in the contiguous United States between June 1^st^ 2006 – May 31^st^ 2018 obtained from the NOAA’s National Climatic Data Center (NCDC; ftp://ftp.ncdc.noaa.gov/pub/data/ghcn/). Length of winter was calculated at each weather station as the number of days with temperatures below 10°C between June 1^st^ and May 31^st^ of the following calendar year. A smooth representation of the length of winter for the contiguous USA (NAD83 projection, resolution of 0.124 degrees) was obtained using anisotropic ordinary kriging performed with ArcGIS (ver 10.7, www.esri.com). The mean winter length was then calculated for each county using the package ‘raster’ (ver 2.9-23) in R (Hijmans 2019 citations). This was the same procedure used in Maher et al. (2012). We used the average length of winter for each county over the full time period as a measure of the winter climate of each county.

In addition to winter length, we also considered the number of days from June 1 until the onset of winter as another indicator of winter conditions, where onset of winter was defined as the first two consecutive days with a minimum daily temperature reading below 10°C for each operational weather station. The number of days until the start of winter for each county was interpolated as above and rounded to the nearest whole number.

### Model

Following Maher et al. (2012), gravity models were fit to the observed time of infection to estimate the effects of weather, distance between counties, cave density, and bat species richness on the spatial spread of the epidemic. All models were of the form

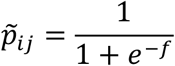

where 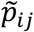 is the probability that county *i* does not become infected from previously infected county *j* and *f* is a function describing the inverse of transmission intensity from *j* to *i*. The basic version of *f* is:

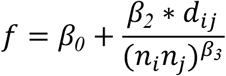

where *d_ij_* is the distance between county *i* and county *j*, *n_i_* is the number of caves in county *i* and *n_j_* is the number of caves in county *j*. The fit parameter *β*_0_ is the background infection rate, and *β*_2_ and *β*_3_ adjust the distance and product of cave densities respectively. Alternative models were constructed by adding additional terms to the function *f* (Supplementary Table 1). We fit a total of 19 models with each winter condition measure crossed with the two species richness estimates.

A change from Maher et al. (2012) is the inclusion of time-varying environmental variables. Previous models used fixed values for all variables representing each county, including the average length of winter in each county over the period of epidemic spread. Here, we also considered models that replaced the average winter conditions with the annual conditions corresponding to each year, as we hypothesized that specific annual conditions might affect either bat behavior or infection intensity and thus influence pathogen transmission (Verant et al., 2012; Langwig et al., 2015; Langwig et al., 2020). We also considered models where the probability of transmission depended on conditions in the prior winter, with the rationale that the observation of WNS in a bat colony (symptomatic disease) may reflect arrival of the fungus (the pathogen) in the previous winter.

Unknown parameters were estimated using maximum likelihood; model selection was performed using Akaike’s Information Criterion (AIC). To consider the role of accumulating information and the influence of unique spread events on model fit, we calculated the AIC value annually for each model. We considered models with consistently low AIC values to be the most robust, but recognize that fluctuations in model performance from year-to-year provide additional insight into the interaction between epidemic pattern and covariates, and guard against model selection being dominated by the conditions of a single, possibly anomalous year at which evaluation occurred. The difference in AIC between the model with the lowest AIC in that year and each of the other models in that year (ΔAIC) was calculated for years 2008-2018. Similarly, we looked at the estimated parameter values for years 2009-2018, focusing on the models that had *Δ*AIC = 0 at some point during the epidemic (Figure 3). We excluded the first two years of estimates because these early estimates were less reliable and exhibited larger changes than for later years (Supplementary Table 2).

## Results

Annual measures of winter severity in U.S. counties were more strongly correlated with average winter severity (Pearson’s r = 0.89) than winter start (Pearson’s r = 0.88, Figure S1). There was noticeable variation from year to year in both measures, as well as signatures of distinct winter patterns in some parts of the United States (Figure 1), for example some areas are always below the temperature that would be considered winter in other areas. Winter length and winter start were strongly negatively correlated, as earlier starts were generally associated with longer winters (Pearson’s r = −0.93, Supplementary Figure 1).

**Figure 1:**
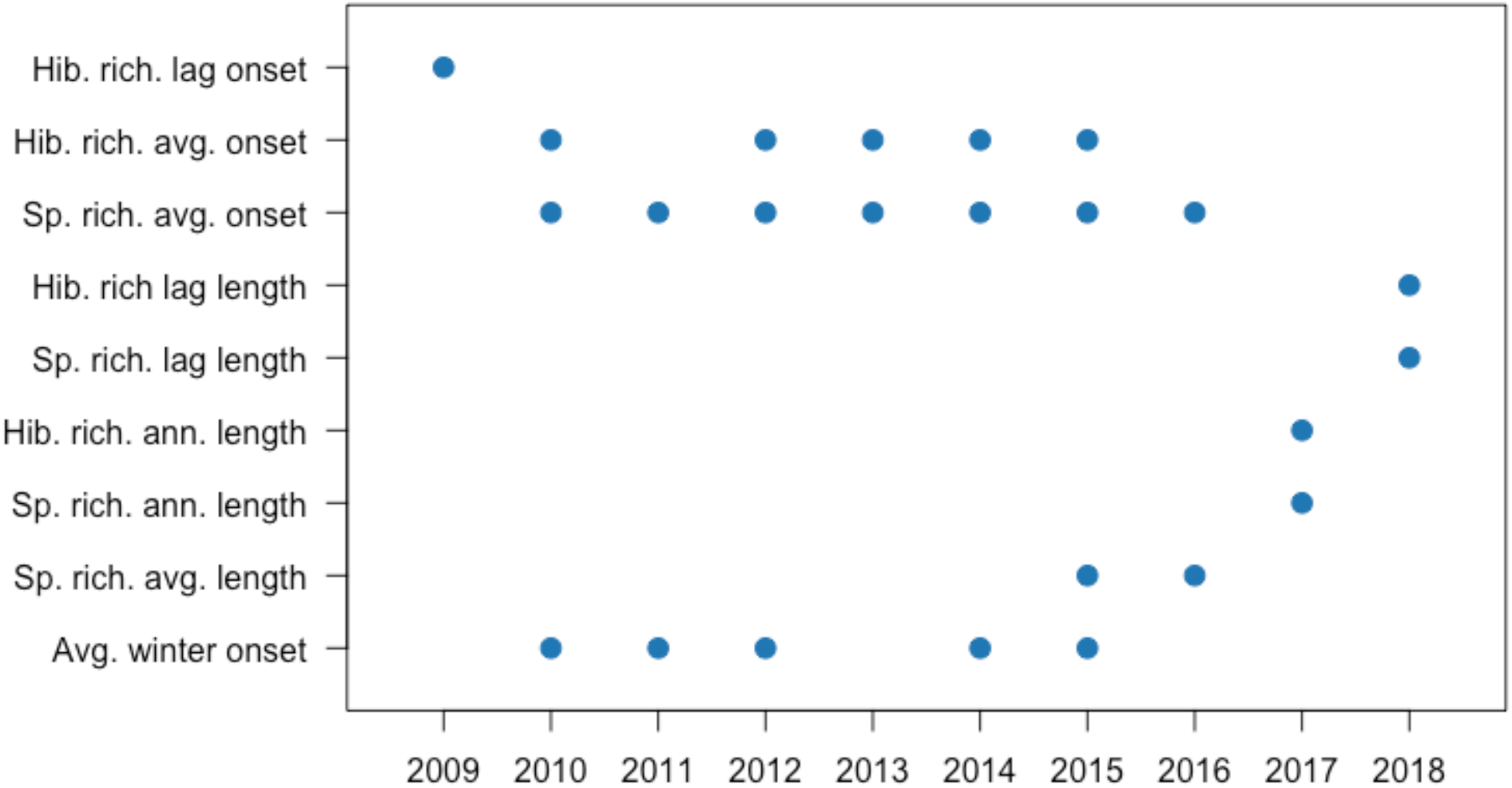
Best-supported models at the end of each year from 2009-2018. All models with *ΔACI* < *2* for any year (from 2009-2018) are included and a mark indicates *ΔACI* < *2* for the given year.

**Figure 2.**
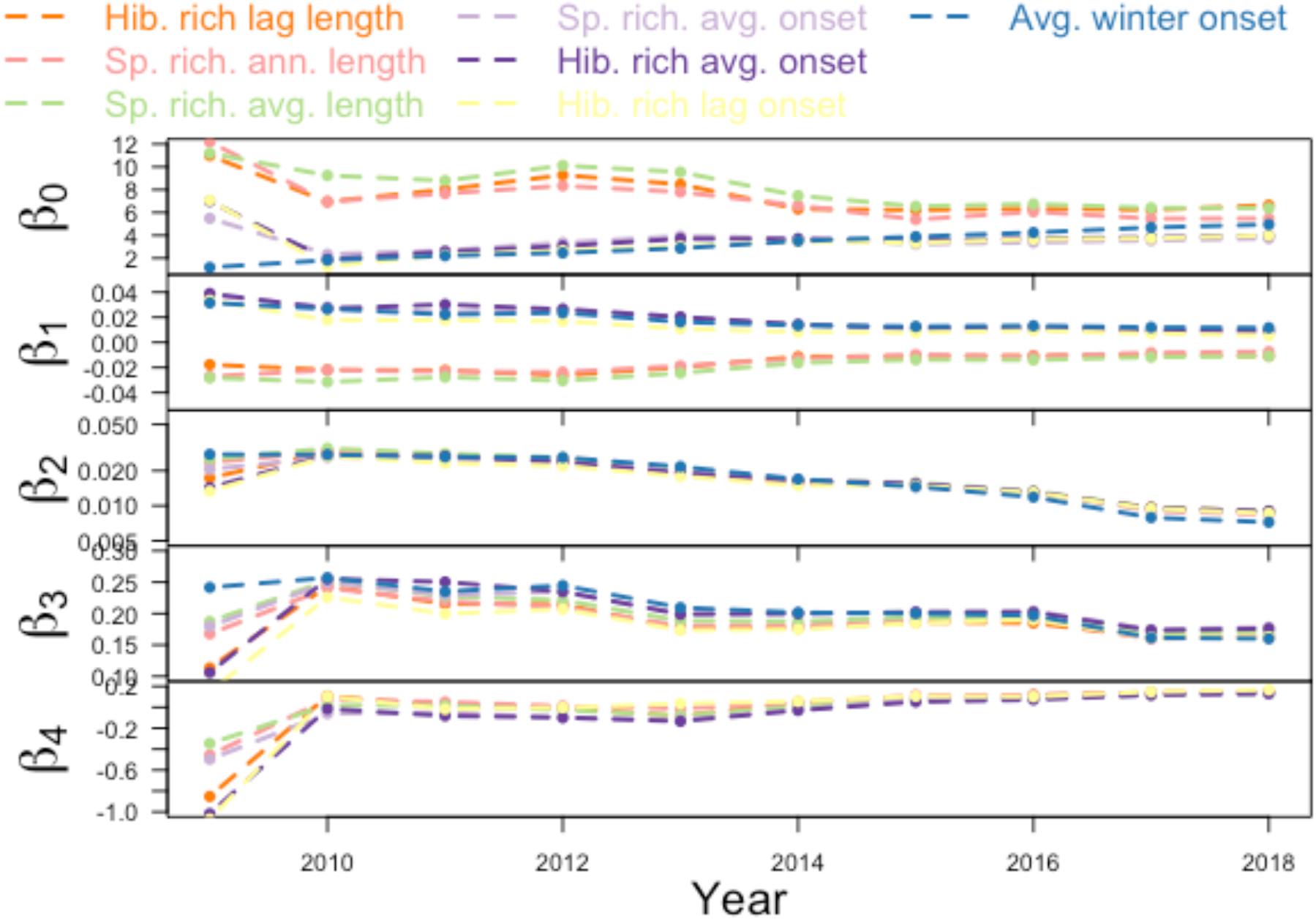
Variation in parameter estimates from models fit at the end of each year from 2009-2018. Selected models appeared in the best fit models and represent several combinations of winter conditions and species richness data.

The best fit model at the end of our study period (winter of 2018-2019) included county-to-county distance, cave abundance, species richness of hibernating bats, and the average length of winter in the previous year (Supplementary Table 1). This model was not the most supported when fit to observations of the WNS outbreak prior to 2018 (Figure 1). Changes in model fit over the course of the epidemic showed that some measure of winter severity was part of the most informative model in every year (Fig 1, Supplementary Table 2). Average measures of winter severity had the lowest AIC in 6 of 11 years, while annual measures from the previous year had the lowest AIC in 3 years and annual measures from the present year had lowest AIC in two years. There were large shifts from year to year in which model was most supported (*Δ*AIC < 2, Figure 1). Species richness was included in 8 out of 9 supported models, including those for all years following the winter of 2014-2015. The start of winter was the most informative measure of winter severity prior to 2015-2016, after which winter length dominated the top performing models. These shifts in the most supported explanatory factors likely indicate some combination of increased discriminatory power as data accumulated and alterations in the spread process as new areas were affected.

Estimated values of parameters associated with background transmission (*β*_0_), winter severity (*β*_1_), distance (*β*_2_), winter severity (*β*_3_), and bat species richness (*β*_4_)varied over the course of the outbreak (Figure 3). Early in the outbreak the models including winter length had a higher estimate of background infection rate, suggesting the other covariates in these models were not capturing as much of the pattern of spread, but in later years the estimates of background transmission were similar across models. (Figure 3a). As expected, winter length was positively correlated with the probability WNS was observed. Similarly, winter start had a negative relationship (Figure 3b), with the magnitude of the effect of winter conditions declining over time (moving towards zero) for both measures. The estimated effect of distance on spread declined by almost an order of magnitude over time in all models (Figure 3c). Cave density had a more stable influence on spread patterns (Figure 3d). Finally, the estimated species richness parameter was stable and similar whether considering total bat richness or hibernating bat richness (Figure 3e).

## Discussion

The well-resolved spatial dynamics of WNS allowed us to distinguish aspects of the dynamics that remained consistent as spread continued across the continent from those that changed as the outbreak proceeded. Winter conditions remained an important factor in spatial dynamics throughout the outbreak, but the most informative measure of winter conditions changed over the course of the epidemic, which could be due to differing drivers in different regions or correlations with unmeasured causal mechanisms related to winter severity. However, our analysis indicates that year-to-year variation in winter conditions is playing a measurable role in epidemic dynamics despite strong correlations with average climate in each county. We found that geographic patterns in species richness are related to the probability of transmission between counties, and detection of this relationship has become more reliable as the outbreak has proceeded. In addition we found that the effect of distance on transmission declined from estimates by Maher et al. (2012), and this decline was gradual, rather than an abrupt response to the 2016 detection of *P. destructans* in Washington State (Lorch et al., 2016).

The long distance transmission event to Washington State was of particular interest as this either represented the realization of an event (albeit one of low probability) encompassed by the model of Maher et al. (2012), or an unforeseen possibility demonstrating that models fit early in the outbreak missed important aspects influencing spread. The suggestion that this spread was human-mediated due to its extreme distance from the nearest known infected hibernacula (Hoyt et al., 2021) could be consistent with either explanation. The inclusion of species richness in most of the supported models in recent years represents one possible signal of the spread to the western United States. The community of bats in the western U.S. has different species and higher richness than the region where WNS emerged, and these areas are separated by a relatively species-poor area through the middle of the country (Maher et al., 2012). At the same time we did not observe an abrupt change in the parameter controlling the influence of distance in the model after this long distance event, instead accumulating data was already indicating distance was less of a barrier than found early in the epidemic.

It is important to note how the geographic pattern in species richness may correlate with additional factors we were unable to consider here. The sign of the estimated parameter means that the model estimated probability of transmission went down as species richness increased. The continental pattern of spread has been from an area of low richness in the Northeast, along the Appalachian Mountains, then towards the Ozark Plateau, where more species ranges overlap. West of the Ozark Plateau, species richness declines into the Great Plains, and then more species occur west of the 100th meridian. As such, the model may reflect a bottleneck of suitable hibernacula that corresponded to a relative increase in richness. Further, the geography of species richness reflects a combination of ecological and evolutionary processes (Miller-Butterworth et al., 2014) and ignores population heterogeneity (e.g. Wilder et al., 2015). Thus, it is possible that the specific mechanism associated with this covariate has nothing to do with the ecological community and it would be premature to conclude that counties with greater species richness in the western U.S. necessarily have low risk of pathogen transmission.

The influence of winter conditions on spatial spread of WNS is of interest not only for understanding relative risk of different bat populations (Maher et al., 2012; Wilder et al., 2011; Lilley et al., 2018), but also because of the importance of environmental conditions in disease severity (Langwig et al., 2012; Hayman et al., 2016) and hypothesized pathways of transmission between hibernacula (Langwig et al., 2020). Research suggests that bat movements during the winter may be a driver of transmission between hibernacula (Langwig et al., 2020) and ambient temperature affects bat activity during winter (Bernard & McCracken, 2017). Our findings suggest that aspects of winter severity related to winter length and the timing of winter onset both improve model performance, with winter onset appearing more informative early in the epidemic, while length of winter is favored over the full period up to the present. We also found evidence that the conditions in particular winters drive transmission risk and that this is detectable despite the strong correlation with average conditions. However, although the previous winter is more valuable for predicting the risk of transmission when looking at the full period to the present, the tendency for the best fit model to vary between years suggests we should not overestimate the strength of this inference. A signal of the previous winter could indicate detection occurring the year after fungal arrival in a hibernacula (Langwig et al., 2015). Improving this aspect of the model may depend on improved understanding of which factors lead to the onset of hibernation or movement among hibernacula.

Examining how models of the spread of WNS have changed over time provided a richer understanding of spatial spread in this ongoing outbreak. At all time points, distance, cave density, and winter conditions best explained the spread, suggesting that even models fit to the exponential phase of an epizootic can provide information sufficient for assessing risk and designing interventions. At the same time, the models best able to explain the observations changed significantly as the outbreak proceeded, as did the relative strength of multiple factors in those models. Hence, the effort of updating the model provides additional insight into WNS dynamics and will facilitate improved forecasts of future spread. This aspect of disease modeling is likely important to any long term outbreak event and highlights the value of revisiting even high performing models.

## Acknowledgements

This research was supported by the NSF Macrosystems Biology program under grant EF-1442417 to Kramer and Drake.

---

**Supplementary Table 1:** Transmission intensity function and AIC by outbreak year for each model (2007 represents infection observed after winter 2006-2007).

| | $f$ | 2007 | 2008 | 2009 | 2010 | 2011 | 2012 | 2013 | 2014 | 2015 | 2016 | 2017 | 2018 |
| --- | --- | --- | --- | --- | --- | --- | --- | --- | --- | --- | --- | --- | --- |
| Gravity (caves) | $\beta_0 + \frac{\beta_2 d_{ij}}{(n_i n_j)^{\beta_3}}$ | 6 | 78 | 255 | 495 | 759 | 1067 | 1455 | 1901 | 2349 | 2751 | 3187 | 3486 |
| Gravity (caves) + average winter length | $\beta_0 + \beta_1 WL_{avg} + \frac{\beta_2 d_{ij}}{(n_i n_j)^{\beta_3}}$ | 8 | 78 | 252 | 472 | 727 | 1010 | 1413 | 1860 | 2290 | 2671 | 3105 | 3403 |
| Gravity (caves) + annual winter length | $\beta_0 + \beta_1 WL_{ann} + \frac{\beta_2 d_{ij}}{(n_i n_j)^{\beta_3}}$ | 8 | 75 | 247 | 475 | 723 | 1008 | 1412 | 1861 | 2302 | 2677 | 3098 | 3407 |
| Gravity (caves) + previous winter length | $\beta_0 + \beta_1 WL_{lag} + \frac{\beta_2 d_{ij}}{(n_i n_j)^{\beta_3}}$ | 8 | 75 | 254 | 475 | 730 | 1007 | 1410 | 1871 | 2303 | 2696 | 3124 | 3367 |
| Gravity (caves) + average winter onset | $\beta_0 + \beta_1 WO_{avg} + \frac{\beta_2 d_{ij}}{(n_i n_j)^{\beta_3}}$ | 8 | 76 | 246 | 466 | 719 | 995 | 1400 | 1849 | 2289 | 2671 | 3108 | 3405 |
| Gravity (caves) + annual winter onset | $\beta_0 + \beta_1 WO_{ann} + \frac{\beta_2 d_{ij}}{(n_i n_j)^{\beta_3}}$ | 8 | 80 | 255 | 475 | 735 | 1057 | 1448 | 1891 | 2323 | 2697 | 3151 | 3434 |
| Gravity (caves) + previous winter onset | $\beta_0 + \beta_1 WO_{lag} + \frac{\beta_2 d_{ij}}{(n_i n_j)^{\beta_3}}$ | 8 | 74 | 240 | 473 | 724 | 1017 | 1424 | 1869 | 2308 | 2682 | 3108 | 3431 |
| Gravity (caves) + average winter length + species richness | $\beta_0 + \beta_1 WL_{avg} + \frac{\beta_2 d_{ij}}{(n_i n_j)^{\beta_3}} + \beta_4 SR$ | 10 | 80 | 252 | 474 | 729 | 1012 | 1413 | 1862 | 2289 | 2665 | 3092 | 3384 |
| Gravity (caves) + average winter length + hibernating species | $\beta_0 + \beta_1 WL_{avg} + \frac{\beta_2 d_{ij}}{(n_i n_j)^{\beta_3}} + \beta_4 HB$ | 10 | 80 | 246 | 474 | 729 | 1011 | 1411 | 1862 | 2290 | 2667 | 3091 | 3382 |
| Gravity (caves) + annual winter length + species richness | $\beta_0 + \beta_1 WL_{ann} + \frac{\beta_2 d_{ij}}{(n_i n_j)^{\beta_3}} + \beta_4 SR$ | 10 | 77 | 246 | 477 | 725 | 1010 | 1414 | 1862 | 2295 | 2667 | 3076 | 3377 |
| Gravity (caves) + annual winter length + hibernating species | $\beta_0 + \beta_1 WL_{ann} + \frac{\beta_2 d_{ij}}{(n_i n_j)^{\beta_3}} + \beta_4 HB$ | 10 | 76 | 239 | 477 | 725 | 1010 | 1413 | 1863 | 2298 | 2671 | 3077 | 3378 |
| Gravity (caves) + previous winter length + species richness | $\beta_0 + \beta_1 WL_{lag} + \frac{\beta_2 d_{ij}}{(n_i n_j)^{\beta_3}} + \beta_4 SR$ | 10 | 77 | 255 | 476 | 732 | 1008 | 1411 | 1871 | 2296 | 2680 | 3103 | 3346 |
| Gravity (caves) + previous winter length + hibernating species | $\beta_0 + \beta_1 WL_{lag} + \frac{\beta_2 d_{ij}}{(n_i n_j)^{\beta_3}} + \beta_4 HB$ | 10 | 77 | 250 | 476 | 732 | 1009 | 1410 | 1872 | 2300 | 2687 | 3105 | 3345 |
| Gravity (caves) + average winter onset + species richness | $\beta_0 + \beta_1 WO_{avg} + \frac{\beta_2 d_{ij}}{(n_i n_j)^{\beta_3}} + \beta_4 SR$ | 10 | 78 | 244 | 468 | 720 | 995 | 1398 | 1850 | 2288 | 2666 | 3096 | 3387 |
| Gravity (caves) + average winter onset + hibernating species | $\beta_0 + \beta_1 WO_{avg} + \frac{\beta_2 d_{ij}}{(n_i n_j)^{\beta_3}} + \beta_4 HB$ | 10 | 78 | 239 | 468 | 723 | 995 | 1397 | 1850 | 2289 | 2669 | 3096 | 3387 |
| Gravity (caves) + annual winter onset + species richness | $\beta_0 + \beta_1 WO_{ann} + \frac{\beta_2 d_{ij}}{(n_i n_j)^{\beta_3}} + \beta_4 SR$ | 10 | 82 | 255 | 477 | 737 | 1045 | 1444 | 1883 | 2307 | 2676 | 3114 | 3399 |
| Gravity (caves) + annual winter onset + hibernating species | $\beta_0 + \beta_1 WO_{ann} + \frac{\beta_2 d_{ij}}{(n_i n_j)^{\beta_3}} + \beta_4 HB$ | 10 | 82 | 251 | 477 | 766 | 1049 | 1448 | 1887 | 2314 | 2684 | 3121 | 3404 |
| Gravity (caves) + previous winter onset + species richness | $\beta_0 + \beta_1 WO_{lag} + \frac{\beta_2 d_{ij}}{(n_i n_j)^{\beta_3}} + \beta_4 SR$ | 10 | 76 | 239 | 475 | 726 | 1019 | 1425 | 1868 | 2298 | 2672 | 3087 | 3396 |
| Gravity (caves) + previous winter onset + hibernating richness | $\beta_0 + \beta_1 WO_{lag} + \frac{\beta_2 d_{ij}}{(n_i n_j)^{\beta_3}} + \beta_4 HB$ | 10 | 76 | 233 | 474 | 726 | 1019 | 1426 | 1869 | 2302 | 2675 | 3089 | 3400 |

**Supplementary Table 2:** *Δ*AIC by outbreak year (2007 represents infection observed after winter 2006-2007)

|  | 2008 | 2009 | 2010 | 2011 | 2012 | 2013 | 2014 | 2015 | 2016 | 2017 | 2018 |
| --- | --- | --- | --- | --- | --- | --- | --- | --- | --- | --- | --- |
| Gravity (caves) | 4.4 | 21.8 | 28.9 | 40.8 | 72.6 | 57.5 | 52.4 | 61.3 | 86.7 | 111.4 | 140.9 |
| Gravity (caves) + average winter length | 3.8 | 18.7 | 5.6 | 8.4 | 15.2 | 15.6 | 11.6 | 2.8 | 5.9 | 29.1 | 58.4 |
| Gravity (caves) + annual winter length | 0.7 | 13.8 | 8.9 | 4.3 | 13.1 | 14.7 | 12.6 | 14.5 | 12.6 | 22.0 | 62.0 |
| Gravity (caves) + previous winter length | 1.1 | 21.2 | 8.5 | 11.4 | 11.9 | 12.3 | 22.6 | 15.4 | 31.7 | 48.5 | 21.8 |
| Gravity (caves) + average winter onset | 2.4 | 12.3 | **0.0** | **0.0** | **0.0** | 3.1 | **0.0** | 1.0 | 6.8 | 32.5 | 60.5 |
| Gravity (caves) + annual winter onset | 6.4 | 21.4 | 9.2 | 16.5 | 62.5 | 50.7 | 42.6 | 35.8 | 32.8 | 74.9 | 89.5 |
| Gravity (caves) + previous winter onset | **0.0** | 6.9 | 7.0 | 5.4 | 22.8 | 27.1 | 20.0 | 20.1 | 17.1 | 32.6 | 86.2 |
| Gravity (caves) + average winter length + species richness | 5.7 | 19.1 | 7.6 | 10.4 | 17.1 | 15.7 | 13.6 | 1.3 | **0.0** | 16.0 | 38.8 |
| Gravity (caves) + average winter length + hibernating species | 5.6 | 12.9 | 7.6 | 10.3 | 16.8 | 14.1 | 13.6 | 2.8 | 2.8 | 15.7 | 37.2 |
| Gravity (caves) + annual winter length + species richness | 2.7 | 13.0 | 10.6 | 6.0 | 15.1 | 16.6 | 13.8 | 7.4 | 2.8 | **0.0** | 32.0 |
| Gravity (caves) + annual winter length + hibernating species | 2.5 | 5.8 | 10.4 | 6.3 | 15.1 | 16.0 | 14.4 | 10.8 | 6.7 | 1.7 | 33.1 |
| Gravity (caves) + previous winter length + species richness | 3.1 | 22.1 | 10.1 | 13.1 | 13.9 | 13.6 | 21.9 | 8.5 | 15.8 | 27.2 | 1.3 |
| Gravity (caves) + previous winter length + hibernating species | 2.8 | 17.0 | 9.9 | 13.4 | 13.9 | 12.2 | 23.4 | 12.4 | 22.5 | 29.3 | **0.0** |
| Gravity (caves) + average winter onset + species richness | 4.4 | 11.0 | 1.8 | 1.7 | 0.2 | 0.8 | 1.7 | **0.0** | 1.7 | 20.0 | 42.4 |
| Gravity (caves) + average winter onset + hibernating species | 4.2 | 5.8 | 2.0 | 4.2 | 0.5 | **0.0** | 1.6 | 1.4 | 4.6 | 20.7 | 42.2 |
| Gravity (caves) + annual winter onset + species richness | 8.2 | 22.2 | 11.2 | 18.0 | 50.8 | 46.9 | 34.2 | 19.5 | 11.7 | 37.8 | 53.9 |
| Gravity (caves) + annual winter onset + hibernating species | 8.3 | 17.3 | 11.1 | 47.4 | 53.9 | 50.3 | 38.5 | 26.7 | 19.9 | 45.3 | 58.7 |
| Gravity (caves) + previous winter onset + species richness | 2.0 | 6.1 | 8.7 | 7.4 | 24.8 | 27.7 | 19.4 | 10.9 | 7.4 | 11.1 | 51.0 |
| Gravity (caves) + previous winter onset + hibernating species | 1.7 | **0.0** | 8.3 | 7.4 | 24.8 | 28.6 | 20.5 | 14.8 | 10.8 | 13.1 | 54.8 |

**Figure S1:**
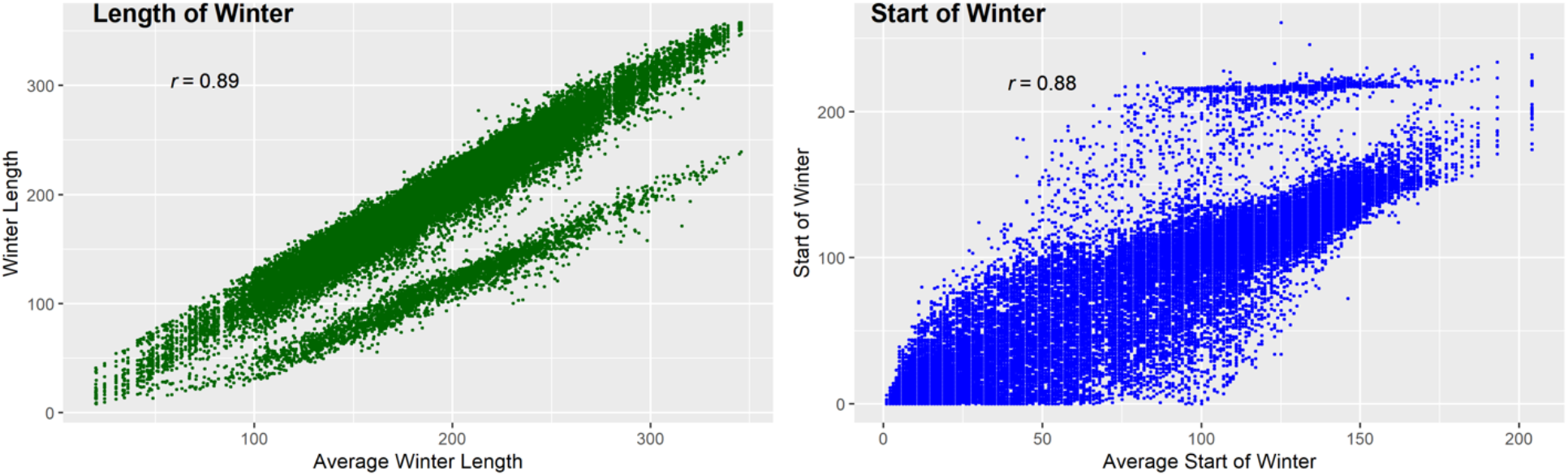
Correlation between average winter conditions and annual winter conditions. A) Winter length for each county over the period June 1, 2005-May 31, 2018. B) Winter onset, as defined by first two consecutive days < 10C, for each county over the period June 1, 2005-May 31, 2018.

**Figure S2:**
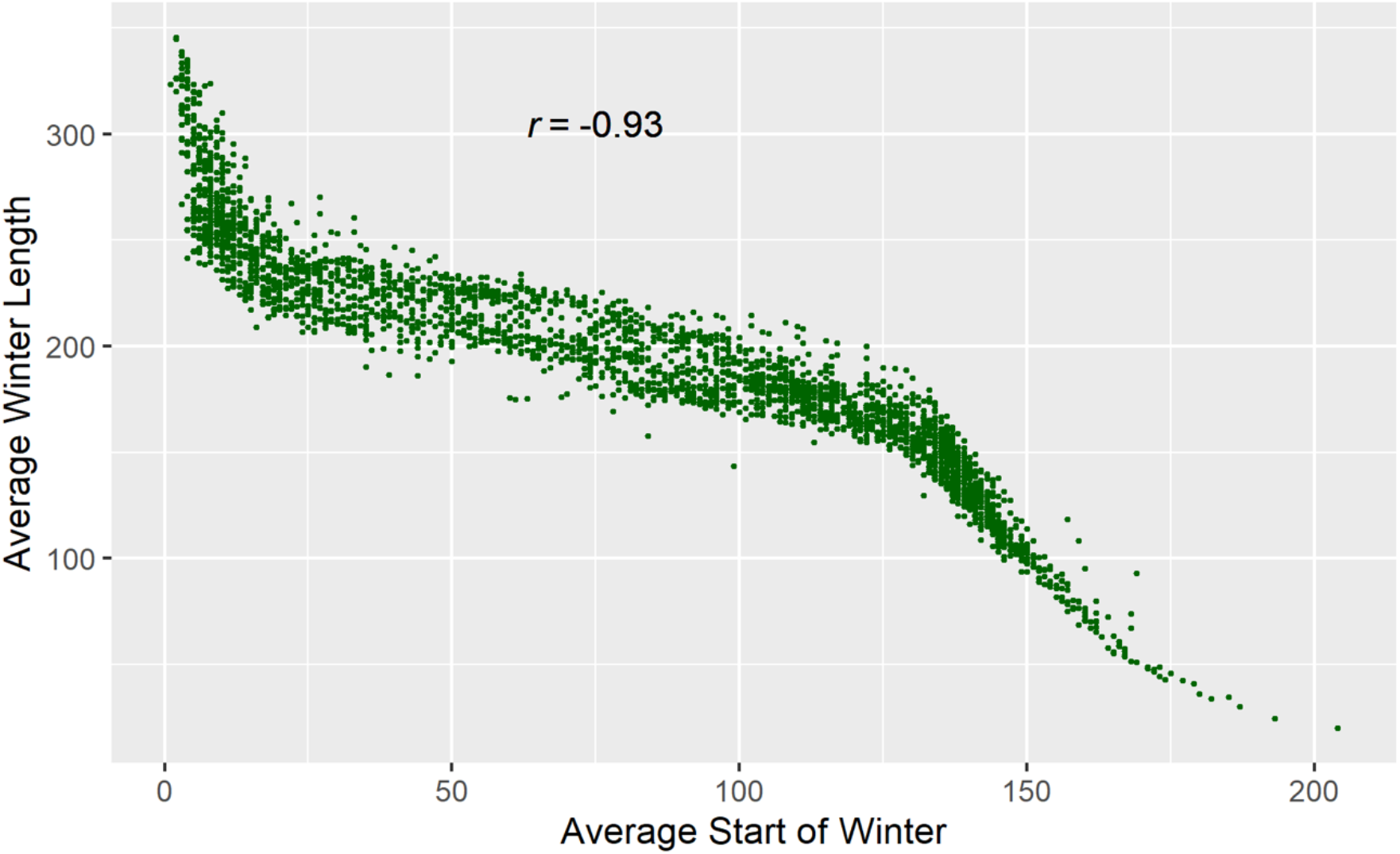
Relationship between average winter length and the average winter onset for U.S. counties.

## Literature Cited

Bernard, R. F., & McCracken, G. F. (2017). Winter behavior of bats and the progression of white-nose syndrome in the southeastern United States. Ecology and Evolution, 7(5), 1487–1496. https://doi.org/10.1002/ece3.2772

Frick, W. F., Pollock, J. F., Hicks, A. C., Langwig, K. E., Reynolds, D. S., Turner, G. G., Butchkoski, C. M., & Kunz, T. H. (2010). An Emerging Disease Causes Regional Population Collapse of a Common North American Bat Species. Science, 329(5992), 679–682. https://doi.org/10.1126/science.1188594

Frick, W. F., Puechmaille, S. J., Hoyt, J. R., Nickel, B. A., Langwig, K. E., Foster, J. T., Barlow, K. E., Bartonička, T., Feller, D., Haarsma, A.-J., Herzog, C., Horáček, I., Kooij, J. van der, Mulkens, B., Petrov, B., Reynolds, R., Rodrigues, L., Stihler, C. W., Turner, G. G., & Kilpatrick, A. M. (2015). Disease alters macroecological patterns of North American bats. Global Ecology and Biogeography, 24(7), 741–749. https://doi.org/10.1111/geb.12290

Hayman, D. T. S., Pulliam, J. R. C., Marshall, J. C., Cryan, P. M., & Webb, C. T. (2016). Environment, host, and fungal traits predict continental-scale white-nose syndrome in bats. Science Advances, 2(1), e1500831. https://doi.org/10.1126/sciadv.1500831

Hijmans, R. J. (2019). raster: Geographic Data Analysis and Modeling. (R package version 2.9-23) [Computer software]. https://CRAN.R-project.org/package=raster

Hoyt, J. R., Kilpatrick, A. M., & Langwig, K. E. (2021). Ecology and impacts of white-nose syndrome on bats. Nature Reviews Microbiology, 1–15. https://doi.org/10.1038/s41579-020-00493-5

Kramer, A. M., Teitelbaum, C. S., Griffin, A., & Drake, J. M. (2019). Multiscale model of regional population decline in little brown bats due to white-nose syndrome. Ecology and Evolution, 9(15), 8639–8651. https://doi.org/10.1002/ece3.5405

Langwig, K. E., Feng, J., Parise, K. L., Kath, J., Kirk, D., Frick, W. F., Foster, J. T., & Kilpatrick, A. M. (n.d.). Invasion Dynamics of White-Nose Syndrome Fungus, Midwestern United States, 2012–2014—Volume 21, Number 6—June 2015—Emerging Infectious Diseases journal—CDC. https://doi.org/10.3201/eid2106.150123

Langwig, K. E., Frick, W. F., Bried, J. T., Hicks, A. C., Kunz, T. H., & Marm Kilpatrick, A. (2012). Sociality, density-dependence and microclimates determine the persistence of populations suffering from a novel fungal disease, white-nose syndrome. Ecology Letters, 15(9), 1050–1057. https://doi.org/10.1111/j.1461-0248.2012.01829.x

Langwig, K. E., Frick, W. F., Hoyt, J. R., Parise, K. L., Drees, K. P., Kunz, T. H., Foster, J. T., & Kilpatrick, A. M. (2016). Drivers of variation in species impacts for a multi-host fungal disease of bats. Philosophical Transactions of the Royal Society B: Biological Sciences, 371(1709), 20150456. https://doi.org/10.1098/rstb.2015.0456

Langwig, K. E., White, J. P., Parise, K. L., Kaarakka, H. M., Redell, J. A., DePue, J. E., Scullon, W. H., Foster, J. T., Kilpatrick, A. M., & Hoyt, J. R. (2020). Mobility and infectiousness in the spatial spread of an emerging fungal pathogen. BioRxiv, 2020.05.07.082651. https://doi.org/10.1101/2020.05.07.082651

Lilley, T. M., Anttila, J., & Ruokolainen, L. (2018). Landscape structure and ecology influence the spread of a bat fungal disease. Functional Ecology, 32(11), 2483–2496. https://doi.org/10.1111/1365-2435.13183

Lorch, J. M., Meteyer, C. U., Behr, M. J., Boyles, J. G., Cryan, P. M., Hicks, A. C., Ballmann, A. E., Coleman, J. T. H., Redell, D. N., Reeder, D. M., & Blehert, D. S. (2011). Experimental infection of bats with Geomyces destructans causes white-nose syndrome. Nature, 480, 376–378. https://doi.org/10.1038/nature10590

Lorch, J. M., Palmer, J. M., Lindner, D. L., Ballmann, A. E., George, K. G., Griffin, K., Knowles, S., Huckabee, J. R., Haman, K. H., Anderson, C. D., Becker, P. A., Buchanan, J. B., Foster, J. T., & Blehert, D. S. (2016). First Detection of Bat White-Nose Syndrome in Western North America. MSphere, 1(4). https://doi.org/10.1128/mSphere.00148-16

Maher, S. P., Kramer, A. M., Pulliam, J. T., Zokan, M. A., Bowden, S. E., Barton, H. D., Magori, K., & Drake, J. M. (2012). Spread of white-nose syndrome on a network regulated by geography and climate. Nature Communications, 3, 1306. https://doi.org/10.1038/ncomms2301

McCallum, H., Tompkins, D. M., Jones, M., Lachish, S., Marvanek, S., Lazenby, B., Hocking, G., Wiersma, J., & Hawkins, C. E. (2007). Distribution and Impacts of Tasmanian Devil Facial Tumor Disease. EcoHealth, 4(3), 318. https://doi.org/10.1007/s10393-007-0118-0

Miller-Butterworth, C. M., Vonhof, M. J., Rosenstern, J., Turner, G. G., & Russell, A. L. (2014). Genetic Structure of Little Brown Bats (Myotis lucifugus) Corresponds with Spread of White-Nose Syndrome among Hibernacula. Journal of Heredity, 105(3), 354–364. https://doi.org/10.1093/jhered/esu012

Rohr, J. R., Raffel, T. R., Romansic, J. M., McCallum, H., & Hudson, P. J. (2008). Evaluating the links between climate, disease spread, and amphibian declines. Proceedings of the National Academy of Sciences, 105(45), 17436–17441. https://doi.org/10.1073/pnas.0806368105

Thogmartin, W. E., King, R. A., McKann, P. C., Szymanski, J. A., & Pruitt, L. (2012). Population-level impact of white-nose syndrome on the endangered Indiana bat. Journal of Mammalogy, 93(4), 1086–1098.

Turner, G. G., Meteyer, C. U., Barton, H., Gumbs, J. F., Reeder, D. M., Overton, B., Bandouchova, H., Bartonička, T., Martínková, N., Pikula, J., Zukal, J., & Blehert, D. S. (2014). Nonlethal screeing of bat-wing skin with the use of ultraviolet fluoresence to detect lesions indicative of white-nose syndrome. Journal of Wildlife Diseases, 50(3), 566–573. https://doi.org/10.7589/2014-03-058

U.S. Geological Survey. (n.d.). White-Nose Syndrome Threatens the Survival of Hibernating Bats in North America. Retrieved January 27, 2021, from https://www.usgs.gov/centers/fort/science/white-nose-syndrome-threatens-survival-hibernating-bats-north-america?qt-science_center_objects=0#qt-science_center_objects

USFWS. (2019). White-Nose Syndrome.http://www.fws.gov/whitenosesyndrome/maps.html

Verant, M. L., Boyles, J. G., Jr, W. W., Wibbelt, G., & Blehert, D. S. (2012). Temperature-Dependent Growth of Geomyces destructans, the Fungus That Causes Bat White-Nose Syndrome. PLOS ONE, 7(9), e46280. https://doi.org/10.1371/journal.pone.0046280

Warnecke, L., Turner, J. M., Bollinger, T. K., Lorch, J. M., Misra, V., Cryan, P. M., Wibbelt, G., Blehert, D. S., & Willis, C. K. (2012). Inoculation of bats with European Geomyces destructans supports the novel pathogen hypothesis for the origin of white-nose syndrome. Proceedings of the National Academy of Sciences, 109(18), 6999–7003.

Wilder, A. P., Frick, W. F., Langwig, K. E., & Kunz, T. H. (2011). Risk factors associated with mortality from white-nose syndrome among hibernating bat colonies. Biology Letters, 7(6), 950–953. https://doi.org/10.1098/rsbl.2011.0355

Wilder, A. P., Kunz, T. H., & Sorenson, M. D. (2015). Population genetic structure of a common host predicts the spread of white-nose syndrome, an emerging infectious disease in bats. Molecular Ecology, 24(22), 5495–5506. https://doi.org/10.1111/mec.13396

